# *Rpt5* encodes a receptor-like protein that provides broad and effective net form net blotch (*Pyrenophora teres* f. *teres*) resistance in barley

**DOI:** 10.1101/2024.08.09.607401

**Authors:** Karl Effertz, Jonathan K. Richards, Shaun J. Clare, Madeline Del Castillo, Roshan Sharma Poudel, Mengyuan Li, Jianwei Zhang, Matthew J. Moscou, Timothy L. Friesen, Robert S. Brueggeman

## Abstract

The foliar disease net form net blotch (NFNB), caused by the necrotrophic fungal pathogen *Pyrenophora teres* f. *teres* (*Ptt*), causes significant yield and quality losses of barley worldwide. Dominant resistance conferred by the <u>R</u>esistance to *<u>P</u>yrenophora teres <u>5</u>*(*Rpt5*) gene from barley line CI5791 is the broadest and most effective resistance reported in this pathosystem. The *Rpt5* locus was identified in multiple independent genetic studies utilizing diverse host populations and *Ptt* isolates, and harbors both dominant *Rpt5* resistance and isolate-specific susceptibility genes/alleles that are dominant in the absence of *Rpt5,* designated <u>s</u>usceptibility to *<u>P</u>yrenophora teres <u>1</u>* (*Spt1*). *Ptt* virulence and avirulence effectors from diverse pathogen isolates genetically interact with the *Rpt5/Spt1* locus, suggesting a complex locus with a function targeted by the evolution of a diversity of pathogen effectors. High-resolution mapping utilizing 1,920 recombinant gametes from a CI5791 x Tifang biparental population, identified 12 candidate genes in an ∼4.6 Mb delimited region in the cv Morex V3 genome assembly, but is 1.1 – 2.2 Mb in the pangenome assemblies, containing 5-12 genes. Analysis revealed a strong correlation between the CI5791 allele of a receptor-like protein (RLP), provisionally designated *<u>R</u>pt5* <u>c</u>andidate <u>g</u>ene <u>1</u>, (*Rcg1*), and broad *Rpt5*-mediated resistance. Two independent transformants of the CI5791 *Rcg1* allele in the susceptible cv Golden Promise background showed significantly increased resistance when challenged with *Rpt5* avirulent *Ptt* isolates 6A, 15A, and 0-1 compared to the Golden Promise wildtype. Thus, *Rpt5,* encodes an RLP and is the first net blotch resistance gene cloned in barley.

**Significance Statement:** Net form net blotch (NFNB) causes substantial yield and quality losses of barley worldwide. Despite decades of ongoing research into genetic resistance to *Pyrenophora teres* f. *teres*, the causal agent of NFNB, no host resistance or susceptibility genes in this pathosystem had been previously identified. We present the identification and validation of *Rpt5*, which confers the broadest and most effective resistance against NFNB. To our knowledge *Rpt5* is the first genetically defined resistance gene in a monocot to encode an RLP.

## Introduction

Fungal diseases are a major yield and quality limiting factor of cereal grain production worldwide. As cereals are a primary source of human caloric intake (Knopff and Morell, 2019), identification, and functional characterization of effective resistance genes to fungal pathogens is important for food security. Intelligent deployment of available genetic resistances is especially important for sustainable crop production in the face of climate change. A major disease affecting yield and quality of barley across growing regions globally is net blotch, caused by the destructive foliar pathogen *Pyrenophora teres*. Net blotch presents in two forms that are phenotypically and genetically distinct; net form net blotch (NFNB) caused by *Pyrenophora teres* f. *teres* (*Ptt*) and spot form net blotch (SFNB) caused by *Pyrenophora teres* f. *maculata* (*Ptm*) (Effertz *et al*., 2021). Despite their genetic and phenotypic differences, several barley resistance loci have been identified that mediate resistance against both forms of this pathogen, including the *Rpt5/Spt1* locus characterized in this study (Clare *et al*., 2020). While predominantly a disease of barley, net blotch infections of wheat fields were recently reported (Garozi *et al*., 2020; Perelló *et al*., 2019; Tóth *et al*., 2008), representing a host jump that could pose a future threat to wheat production.

Research investigating genetic resistance to *P. teres* has been ongoing since 1928 (Geschele, 1928). Early studies suggested a single dominant gene was responsible for effective resistance (Geschele, 1928), however subsequent research using genetically diverse host populations identified dominant, recessive, and incomplete dominant resistances (Cakir *et al*., 2003; Khan and Boyd, 1969; Ma *et al*., 2004; Steffenson *et al*., 1996). A locus on barley chromosome 6H has been repeatedly identified as the broadest and most robust source of genetic resistance to *Ptt*, harboring the resistance gene <u>R</u>esistance to *<u>P</u>yrenophora teres* 5 (*Rpt5)* (Manninen *et al*., 2006). The most effective resistances are contributed by the *Rpt5* allele present in Ethiopian landraces CIho5791 (hereafter referred to as CI5791) and CIho9819 (hereafter referred to as CI9819) (Koladia *et al*., 2017; Manninen *et al*., 2006; Robinson and Jalli, 1996). Isolate specific *Ptt* dominant susceptibility designated as <u>S</u>usceptibility to *<u>P</u>yrenophora teres <u>1</u>* (*Spt1*) (Abu Qamar *et al*., 2008; Richards *et al*., 2016) has also been mapped to this locus, as well as *Ptm* resistance (Daba *et al*., 2019; Grewal *et al*., 2012; Tamang *et al*., 2019). Diverse *Ptt* virulence and avirulence effectors (Koladia *et al*., 2017; Liu *et al*., 2015; Richards *et al*., 2016; Shjerve *et al*., 2014) have been shown to interact with the *Rpt5/Spt1* locus in an isolate specific manner. This demonstrates complex genetic interactions driven by strong selection pressure applied on the *Rpt5/Spt1* locus by the pathogen as well as by this host locus on pathogen populations, indicating a complex evolutionary history of the host-pathogen molecular arms race (Jones and Dangl, 2006). Despite the *Rpt5*/*Spt1* locus being identified using diverse host and pathogen populations reported in over 20 publications (Abu Qamar *et al*., 2008; Adhikari *et al*., 2020; Amezrou *et al*., 2018; Daba *et al*., 2019; Emebiri *et al*., 2005; Graner *et al*., 1996; Grewal *et al*., 2008; Gyawali *et al*., 2019; Khan and Boyd, 1969; Ma *et al*., 2004; Manninen *et al*., 2000, 2006; O’Boyle *et al*., 2014; Raman *et al*., 2003; Richards *et al*., 2016, 2017; Vatter *et al*., 2017; Wang *et al*., 2015; Wonneberger *et al*., 2017), efforts have been unsuccessful in the identification of *Rpt5* or *Spt1* or any *P. teres* resistance or susceptibility genes. The mapping of *Rpt5* and *Spt1* to the same genomic location, in addition to the diverse pathogen effectors interacting genetically with this region, suggest that there could potentially be multiple genes involved in the barley - *P. teres* pathosystem underlying this locus.

Quantitative trait loci (QTL) mapping of *Rpt5* was originally published by Manninen et al. (2006), utilizing a doubled haploid (DH) population of 119 individuals resulting from a cross of Finnish 6-row barley Rolfi (susceptible) and Ethiopian landrace CI9819 (resistant). This study found that this resistance was effective against four *Ptt* isolates originating from the US, UK, Canada, and Australia (Manninen *et al*., 2006). Further QTL mapping of the *Rpt5* locus was published in Koladia et al. (2017), using a population of 117 recombinant inbred lines (RILs) generated from a CI5791 x Tifang cross phenotyped with nine geographically diverse *Ptt* isolates. Two major resistance QTL were identified on chromosomes 3H and 6H, respectively. The 6H resistance locus contributed by line CI5791 localized to *Rpt5* and conferred resistance to all nine diverse *Ptt* isolates (V. M. Koladia *et al*., 2017).

We performed high-resolution mapping of *Rpt5*, delimiting the locus to an ∼4.6 Mb region containing 12 genes in the Morex V3 assembly (Figure 1) with four strong candidate *Rpt5* genes provisionally designated *Rpt5* candidate genes 1-4 (*Rcg1-4*). *Rcg1* was predicted to encode a receptor-like protein (RLP) and comparative allele analysis between CI5791 (resistant) and Tifang (susceptible) revealed remarkable diversity. Additionally, allele analysis and disease phenotyping from 80 diverse barley genotypes showed that 100% of individuals (39/39) harboring the CI5791-like allele were highly resistant to all tested *Rpt5* avirulent *Ptt* isolates (6A, 15A, TD10, LDN89-19, and/or 0-1). Stable transformation of the CI5791 *Rcg1* allele driven by a *Mla6* promoter in the susceptible barley cultivar (cv) Golden Promise background resulted in significant shifts towards resistance in two independent transgenic lines. These results show that the RLP is *Rpt5*, the most effective NFNB resistance gene in barley, representing the first *Pyrenophora teres* resistance gene identified and the first genetically defined RLP encoding resistance gene in a monocot.

**Figure 1.**
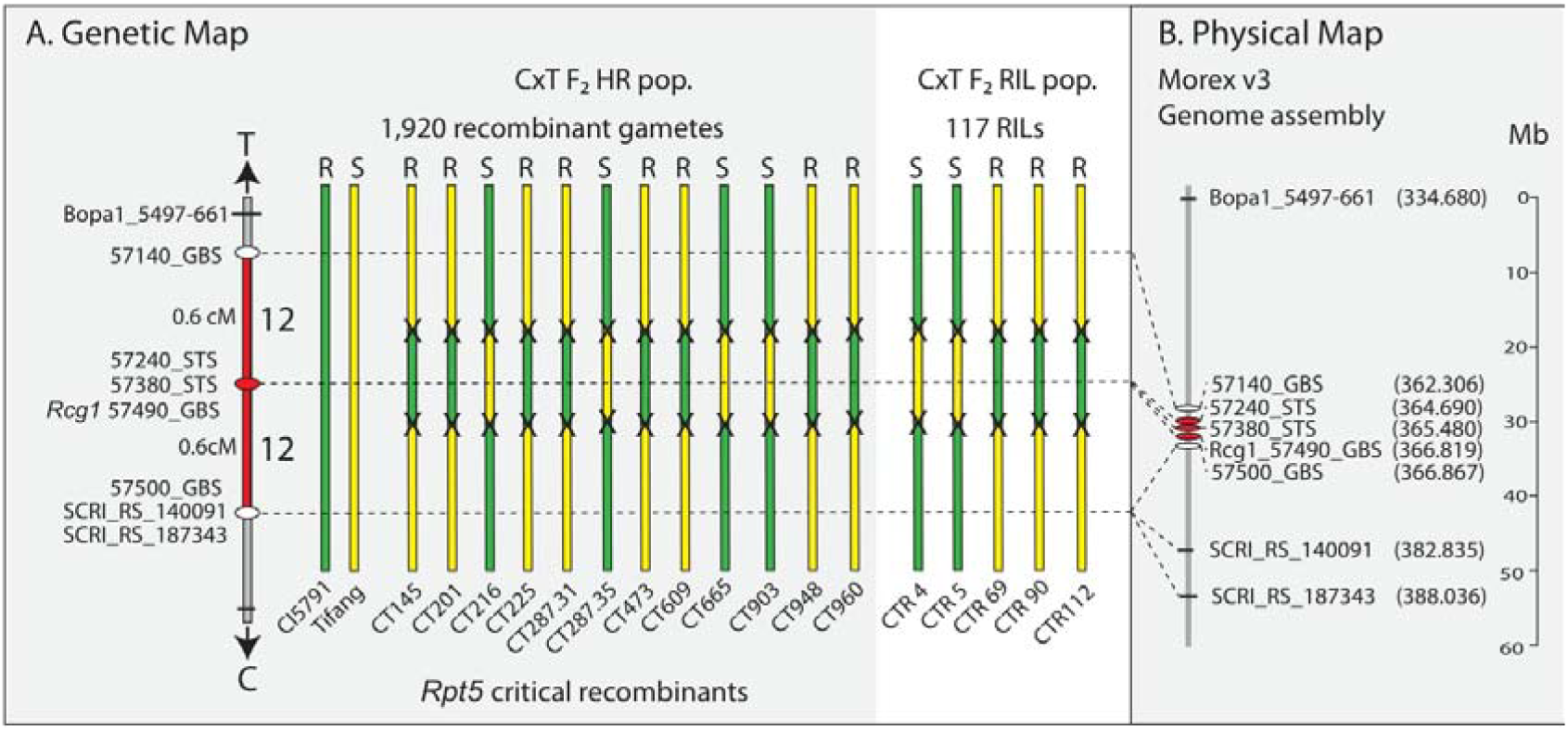
High resolution mapping of *Rpt5*. Genetic (**A**) and physical (**B**) map showing the results of CI5791 x Tifang high-resolution mapping (left). Approximately 1% of F_2_ individuals screened harbored a double recombination surrounding the *Rpt5/Spt1* locus. Double recombination events were originally detected with the Rcg1_57490_GBS marker and confirmed with 57240_STS and 57380_STS markers which correlated perfectly with disease outcomes. STS markers were utilized to screen a CI5791 x Tifang RIL population (right) and a Steptoe x Morex DH population (shown in Dataset S1), confirming that approximately 3-4% of individuals from each of these populations also harbored these double recombination events.

## Results

### High resolution mapping and candidate gene identification

The barley cv Morex V1 genome assembly (Mascher *et al*., 2017) was utilized to design SNP markers for use on the Ion Torrent sequencing platform (Richards *et al*., 2016) targeting the delimited *Rpt5* genic region including the proximal and distal flanking markers 11_20946 and SCRI_RS_140091 (Comadran *et al*., 2012; V. M. Koladia *et al*., 2017), respectively. Using 960 CI5791 x Tifang F_2_ individuals, representing 1,920 recombinant gametes, the delimited *Rpt5* region was saturated with molecular markers. The genotyped individuals were phenotyped with *Ptt* isolate 0-1. Individuals harboring recombination at the *Rpt5* locus were allowed to self and a single F_2:3_ individual homozygous for the recombinant gametes were selected representing 12 immortal critical recombinant lines (Figure 1). In approximately one percent of F_2_ individuals screened, a non-canonical double recombination event occurred delimiting *Rpt5* to a 1.3 cM interval representing ∼4.6 Mb of physical genome sequence from the Morex V3 genome assembly (Figure 1). A single GBS marker, 57490_GBS (within gene *HORVU6Hr1G057490*, Morex V3 ID = *HORVU.MOREX.r3.6HG0595020*), located at 366,819,681 Mb on chromosome 6H correlated perfectly with disease outcome and enabled the detection of these double recombination events between markers 57140_GBS (at 362,306,750 Mb) and 57500_GBS (at 366,867,734 Mb). Additional STS markers were developed targeting the genes *HORVU6Hr1G057240* (Morex V3 ID = *HORVU.MOREX.r3.6HG0594730*) and *HORVU6Hr1G057380* (Morex V3 ID = *HORVU.MOREX.r3.6HG0594810*) within the *Rpt5* region delimited by the double recombinations, confirming these events, and providing additional markers perfectly correlated with disease outcomes (Figure 1). The delimited region was predicted to contain 12 high confidence genes within the Morex V3 assembly (all annotations in Table S3). We employed low coverage PacBio sequencing of resistant parent CI5791 for allele analysis to determine the top candidate gene(s). STS markers and known gene sequences within the delimited region from susceptible parent Tifang were found to be 100% identical to reference cv Morex. This enabled allele comparisons between Morex (as a stand-in for parent Tifang) and CI5791. This analysis revealed that ten genes within the delimited region harbored primary non-conservative amino acid polymorphism between Morex and CI5791. Subsequent analysis utilizing the recently published barley pangenome (Jayakodi *et al*., 2020) revealed that the locus is 1.1 - 2.2 Mb in the majority (17/20) of pangenome lines, harboring 5-12 genes. Of these 5-12 annotated genes within the pangenome, four were homologous to genes previously implicated in plant immune responses designated *<u>R</u>pt5* <u>c</u>andidate <u>g</u>ene <u>1</u> (*Rcg1 - HORVU.MOREX.r3.6HG0595020)* and *Rcg4* (*HORVU.MOREX.r3.6HG0594600*) predicted to encode receptor-like proteins, and *Rcg2 (HORVU.MOREX.r3.6HG0595000)* and *Rcg3 (HORVU.MOREX.r3.6HG0594730)* that encode predicted receptor-like kinases (RLKs). Presence/absence of candidate *‘Rcg’* genes varies greatly across pangenome lines, with only Morex and Golden Promise harboring all four within the *Rpt5*-delimited region. While both *Rcg1* and *Rcg4* are predicted to encode RLPs, they only share ∼71% identity in Morex. Likewise, *Rcg2* and *Rcg3*, both predicted to encode RLKs only share ∼62% identity in Morex. BLAST searches of the CI5791 assembly generated in this study using the Morex *Rcg4* allele suggest that this gene may not be present in CI5791, as the top hit maps to the CI5791 *Rcg1* allele with lower identity and coverage than the Morex *Rcg1* allele. Highly diverged copies of *Rcg2* (∼62% identity) and *Rcg3* (∼73% identity) do appear to be present in the CI5791 genomic sequence, however multiple attempts to amplify full length alleles of *Rcg2* and *Rcg3* from CI5791 cDNA have failed. Investigations into the potential function of *Rcg2* and *Rcg3* in *Rpt5*- mediated resistance or *Spt1*-mediated susceptibility are ongoing, as multiple genes may contribute to the interactions present at this locus. Interestingly, the nearest pangenome relative to CI5791 at this locus (cv RGT Planet) does not contain *Rcg2*, *Rcg3,* or *Rcg4* within the *Rpt5* delimited region. RGT Planet contains seven total high confidence genes within the *Rpt5* locus, with only three showing primary amino acid diversity when compared to CI5791, including *Rcg1*.

### Conserved double recombination events delimit *Rpt5* in multiple populations

High-resolution mapping with the CI5791 x Tifang F_2_ population identified double recombination events delimiting the *Rpt5* locus in approximately one percent of the recombinant gametes with no progeny containing recombination events within the region (Figure 1). To further investigate this phenomenon, STS markers co-segregating with *Rpt5* in the high-resolution mapping were utilized to analyze two additional populations, a CI5791 x Tifang RIL population (Koladia *et al*., 2017) and a Steptoe x Morex DH population (Chen and Hayes, 1989) for the presence of similar double recombination event. These populations were previously genotyped with the iSelect 9k SNP chip and phenotyped with *Ptt* (Koladia *et al*., 2017; Steffenson *et al*., 1996). Our analysis showed that the iSelect 9k SNP chip did not contain polymorphic markers within the double recombination region. The previous mapping of *Ptt* resistance in these populations identified individuals with genotypes flanking the double recombination region that did not correlate with phenotype. These individuals were further genotyped with additional markers within the *Rpt5* region and both the CI5791 x Tifang RIL population (Koladia *et al*., 2017) and Steptoe x Morex DH population (Chen and Hayes, 1989) contained approximately four percent of recombinant progeny harboring double recombination events with similar proximal and distal recombination sites to our high resolution CI5791 x Tifang population (Figure 1). The genotypes of the STS markers within the double recombination region perfectly correlated with disease outcome, explaining the discrepancies in genotype and phenotype correlation for these individual recombinant progeny in the previous genetic mapping (Koladia *et al*., 2017; Steffenson *et al*., 1996). Thus, the double recombination occurs at the *Rpt5*/*Spt1* locus at a relatively high frequency in multiple independent bi-parental populations.

### Predicted structure and cloning of *Rpt5*

Protein domain and structure predictions of *Rcg1* alleles were performed using the InterProScan (Blum *et al*., 2021), Pfam (Mistry *et al*., 2021), DeepTMHMM (Hallgren *et al*., 2022; Kahsay *et al*., 2005), and AlphaFold2 web interfaces (Jumper *et al*., 2021; Varadi *et al*., 2022). These analyses of *Rcg1* revealed typical RLP structures comprised of an N-terminal signal peptide and transmembrane domain, and C-terminal leucine rich repeats (LRRs) predicted as an extracellular receptor domain (Figure 2). Comparative analysis between the CI5791 and Tifang allele sequences showed that *Rcg1* has a high level of diversity (68% amino acid (AA) identity and 76% AA similarity) with a semi-conserved C-terminal leucine rich repeat (LRR) region, a hypervariable hinge region near the proximal end of the predicted LRR domain, a predicted transmembrane domain, and a less conserved N-terminal region harboring a signal peptide (Figure 2). Alignment of predicted AA sequences of CI5791 and Tifang revealed a high level of polymorphism in the extracellular LRR domain. Pfam predicts two LRR4 (PF12799) repeats and three LRR8 (PF13855) repeats within the LRR domain of the CI5791 *Rcg1* allele while Tifang is predicted to harbor a single full-length LRR4 repeat, one truncated LRR4 repeat, and two LRR8 repeats within its predicted LRR domain. Further analysis utilizing the AlphaFold2 predicted structures of CI5791 and Tifang RCG1-extracellular regions revealed differences in the predicted ligand binding pocket. Both isoforms are predicted to contain nine LRR motifs within the ectodomain, each with a hypervariable hinge region in between specific LRRs. In CI5791 this hinge region is predicted between the third and fourth LRR motifs, while in Tifang this hinge is predicted to be located between the fourth and fifth LRR motifs (Figure 3). These differences in LRR organization and binding pocket structure could impact ligand binding affinity, possibly enabling the CI5791 RCG1 receptor to recognize different ligands in the apoplast, which could potentially function in early pathogen recognition, defense signaling and resistance. Annotated *Rcg1* alleles in the barley pangenome range from 1,278 to 1,431 nucleotides, each containing one to two exons, encoding 425 to 476 amino acid proteins. Cultivar RGT Planet harbors the most similar *Rcg1* allele to CI5791 in the pangenome, with 84% nucleotide identity and 78% amino acid identity, demonstrating the extreme variability of this gene. Due to the known function of RLPs as plant immunity receptors (Tang *et al*., 2017) and the predicted domain differences between CI5791 and Tifang alleles, particularly within the extracellular LRR region, *Rcg1* was considered our strongest *Rpt5* candidate gene.

**Figure 2.**
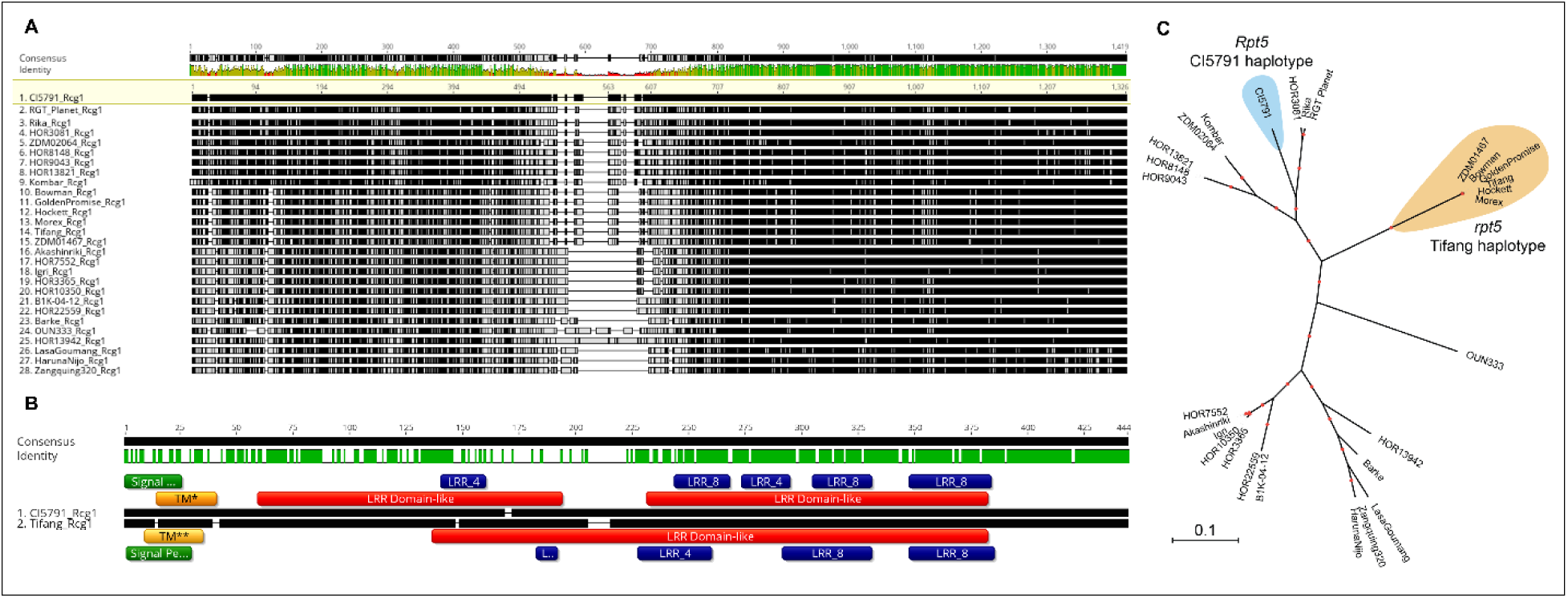
*Rcg1* gene family exhibits extreme diversity. Figure 2. A) Nucleotide alignment of known *Rcg1* alleles including pangenome lines. *Rcg1* alleles are remarkably diverse across barley genotypes, with the greatest conservation observed in the 3’ end of the gene, covering some of the predicted LRR domains. Alignment was performed using the Clustal Omega alignment plugin in Geneious v2022.2.2. B) Peptide alignment of Rcg1 isoforms from resistant parent CI5791 and susceptible parent Tifang. Multiple differences are observed between the specific LRR domains in the predicted extracellular region, which could potentially result in the binding of alternate ligands. C) Maximum likelihood phylogenetic tree of the *Rcg1* gene family based on translation alignment using 31 barley accessions (not shown are Rabat071, Conlon, and CI9819, which are identical to CI5791). Branch support greater than 80% is shown as red dots on branches based on 1,000 bootstraps. The shaded blue region shows the resistant *Rpt5* haplotype of CI5791, while the region shaded in orange is the susceptible rpt5 haplotype of mapping parent Tifang, which correlates with susceptibility to *Ptt* isolates 0-1/TD10.

**Figure 3.**
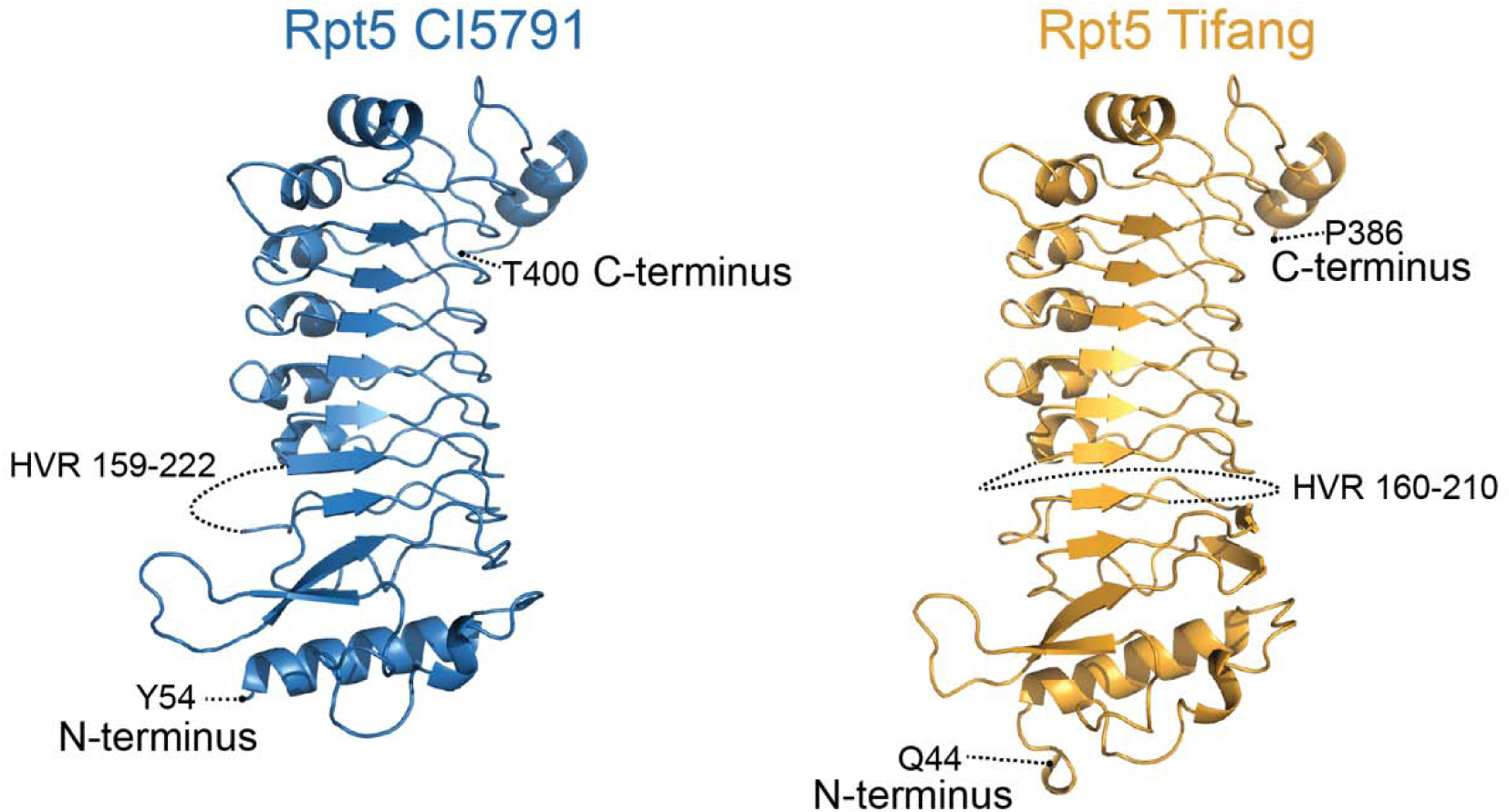
Structure prediction of the ectodomain of Rpt5. Protein structure was predicted using AlphaFold2 with default parameters. Signal peptide/transmembrane region, hypervariable region (HVR), and C-terminal regions were removed from predicted structures to show conservation of leucine-rich repeats. Positions of these regions are indicated.

### Analysis of *Rcg1* alleles from diverse barley cultivars

The high level of sequence diversity observed in *Rcg1* alleles of mapping parents CI5791 and Tifang led us to examine the allelic diversity present in other barley genotypes. We employed a pooled primer system to amplify *Rcg1* alleles from a diverse group of world barley core collection lines (Muñoz-Amatriaín *et al*., 2014) and cultivars that harbored differential reactions to various *Ptt* isolates. These lines had been previously phenotyped with multiple *Rpt5* avirulent *Ptt* isolates (Richards *et al*., 2017). *Rcg1* alleles were amplified with the pooled primers (outlined in Table S2), gel purified, and sequenced on the Ion Torrent PGM sequencing platform. Analysis of the assembled allele sequences revealed four predominant haplotypes from the 80 samples, R1-R4. Interestingly, all lines (n = 39) that possessed the CI5791-like allele of *Rcg1* (R1) were resistant to all isolates tested, while the majority of lines possessing an alternate *Rcg1* allele displayed isolate specific susceptible interactions with at least one of the tested isolates presumably due to the presence of *Spt1* alleles that impart specific susceptibilities (Abu Qamar *et al*., 2008; Richards *et al*., 2016, 2017). Further analysis of the recently released barley pangenome (Jayakodi *et al*., 2020) revealed that no pangenome lines harbor the CI5791 *Rcg1* allele, and consequently all pangenome lines phenotyped were susceptible to at least one of the isolates tested (Table S1). We also phenotyped Ethiopian landrace CI9819 with these isolates, which is known to harbor broad *Ptt* resistance conferred by *Rpt5* (Manninen *et al*., 2006, 2000; Robinson and Jalli, 1996). CI9819 was found to be highly resistant to all *Ptt* isolates tested (average disease score ∼1.84), and sequencing of *Rcg1* from CI9819 showed 100% identity to the CI5791 *Rcg1* allele further demonstrating the efficacy of this allele in conferring resistance to *Ptt*.

### Transformation, phenotyping, and expression analysis of transgenic lines

The full-length CI5791 *Rcg1* allele was cloned into barley transformation vector pBract202 with an *Mla6* promoter, 5’UTR, 3’UTR, and terminator (*Mla6*: *Rcg1.CI5791)*. The *Mla6* promoter and terminator were utilized due to the lack of available CI5791 promoter or terminator sequences. Barley transformations were performed using Golden Promise, which is amenable to *Agrobacterium*-mediated immature embryo stable transformation (Tingay *et al*., 1997) and susceptible to nearly all *Ptt* isolates including *Ptt* isolates 0-1, 6A and 15A. Moreover, Golden Promise harbors a *Rcg1* allele that is highly diverse from CI5791, making it an ideal cultivar for *Rpt5*-validation. Immature barley embryos were transformed resulting in 19 calli and 48 mature transformants. Transgene integration was confirmed via qPCR (Collier *et al*., 2017) with transformants harboring between one and five copies of the Mla6:*Rcg1.CI5791* insert.

Initial phenotypic screening was performed with *Ptt* isolate 6A as described previously at Washington State University, resulting in the identification of two single copy insertion lines (HVT02681 and HVT02690) that showed a shift from an average disease score of 8.1 in Golden Promise wildtype, to average scores of 4.1 and 4.3 for HVT02681 and HVT02690, respectively. Independent phenotyping of the transgenic lines was also performed at the Northern Cereal Crops Research facility (Edward T. Schafer Agricultural Research Center, Fargo, ND) (Friesen *et al*., 2006). This independent phenotyping of the transgenic lines HVT02681 and HVT02690 confirmed the shift from susceptibility to resistance with average scores of 3.2 and 3.5, respectively, compared to average score of 5.7 for Golden Promise when challenged with *Ptt* isolate 6A. Additional phenotyping was carried out for these lines with *Ptt* isolates 0-1, 15A, and TD10. The lowest phenotypic shift was observed with isolate TD-10, with Golden Promise scoring 9.3, HVT02681 scoring 8.2, and HVT02690 scoring 7.5. When challenged with isolate 0-1, Golden Promise displayed a disease reaction of 9.25, with HVT02681 and HVT02690 showing reactions of 7.0 and 7.5, respectively. Phenotyping these lines with isolate 15A revealed a reaction similar to that observed with isolate 6A, with Golden Promise averaging a score of 6.6, HVT02681 averaging 3.1 and HVT02690 averaging 2.8. Additionally, all lines obtained from the same calli as the resistant transgenics (HVT02681, HVT02682 – callus #16, HVT02690, HVT02691, HVT02693 – callus #19) were phenotyped with isolates 0-1, 6A, and 15A, and displayed similar shifts to those noted above for HVT02681 and HVT02690. Statistical analysis utilizing pairwise student’s t-tests revealed that all lines originating from callus #16 and callus #19 were significantly more resistant (*p* < 0.001) than Golden Promise when challenged with all isolates except for TD10. Transgenic lines showed a small numeric shift towards resistance with TD10; however, this shift was not significant (*p* > 0.05) with all transgenic lines evaluated. All transgenic phenotyping results are outlined in Table 1 and Figure 4. A pattern emerged that *Ptt* isolates that were more virulent on parental line Golden Promise displayed the lowest shift in phenotype in our transgenic lines. We predict this is due to additional susceptibility factors that are present in Golden Promise that limit the efficacy of *Rpt5*-mediated resistance, as the phenotyping of F_2_ individuals from a CI5791 x Golden Promise cross did not result in a ratio of 3:1 R:S when challenged with *Ptt* isolate 0-1 suggesting multiple factors contribute to disease phenotype in this cross (Figure S1).

**Figure 4.**
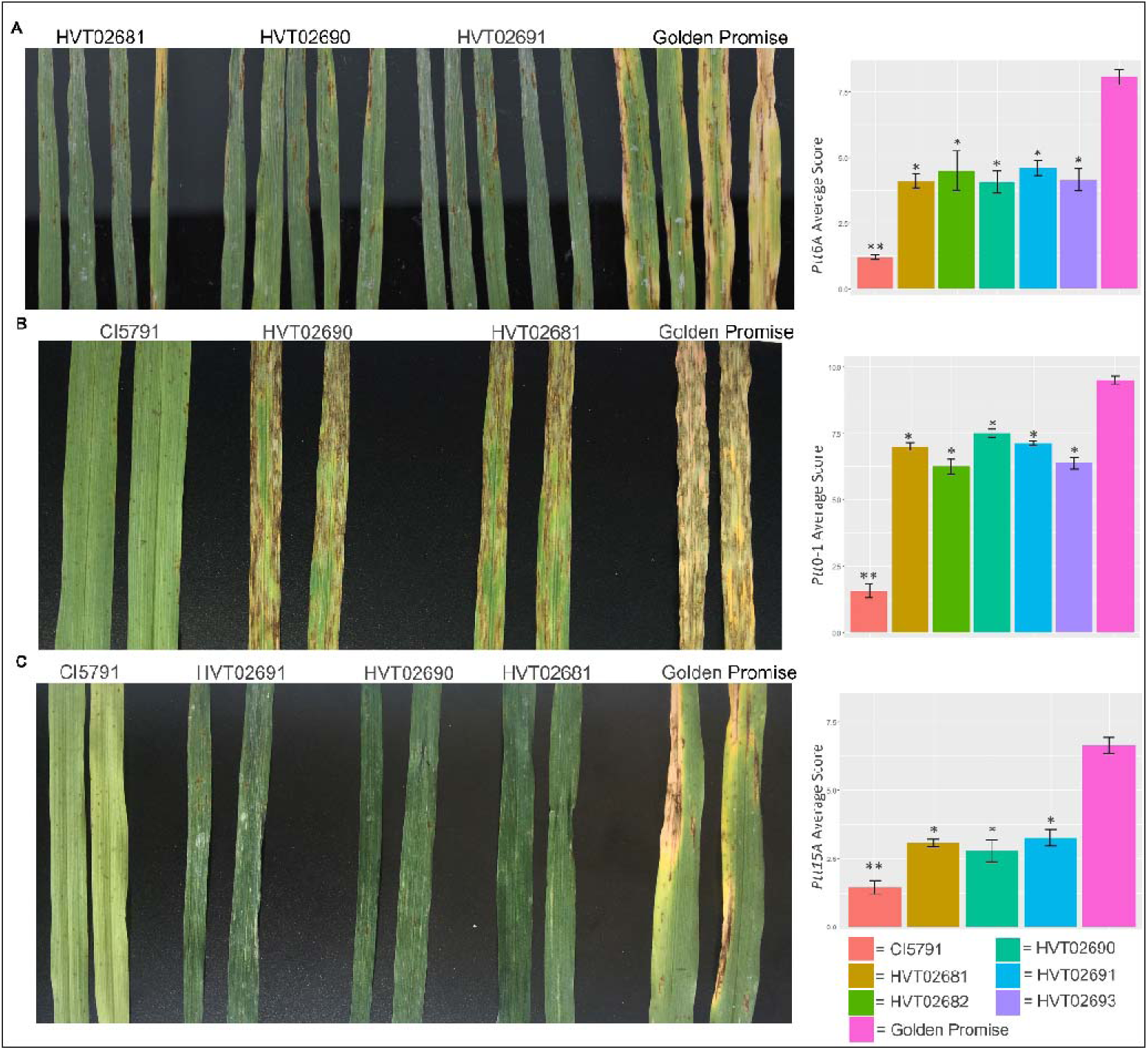
*Rpt5.CI5791* transgenics lines show improved resistance to diverse *Ptt* isolates. Leaf photographs showing Golden Promise and transgenic lines one-week post-inoculation with *Ptt* isolates 6A (**A**), 0-1 (**B**) and 15A (**C**). Bar graphs showing the average disease score for each line are shown on the right of each leaf photo, error bars represent +/- standard deviation. All transgenics originating from callus #16 (HVT02681, HVT02682) and callus #19 (HVT02690, HVT02691, HVT02693) showed a significant shift towards resistance.. None of the transgenic lines showed complete resistance (pinpoint lesions) as observed for CI5791 the source of *Rpt5*, pictured on the left in panels (**B**) and (**C**).

**Table 1.** Summary of parental and transgenic disease *Ptt* disease reactions. Inoculations were performed as described in Friesen *et al*. (2006) and plants were rated using the Tekauz 1-10 rating scale. All averages are based off of at least three biological replications, except those in red, which only had two replications due to low seed number and poor germination. Pairwise student’s t-tests were performed between all lines and Golden Promise to determine statistical significance (* = *p* < 0.05, ** = *p* < 0.001, ns = not significant). Standard deviation is reported on the +/- lines, “np” indicates no plant present for phenotyping.

| WSU Phenotyping | <u>CI5791</u> | <u>HVT02681</u> | <u>HVT02682</u> | <u>HVT02690</u> | <u>HVT02691</u> | <u>HVT02693</u> | <u>Golden Promise</u> |
| --- | --- | --- | --- | --- | --- | --- | --- |
| 0-1 Average | 1.56<sup>**</sup> | 7.00<sup>**</sup> | 6.25<sup>**</sup> | 7.50<sup>**</sup> | 7.13<sup>**</sup> | 6.38<sup>**</sup> | 9.50 |
| +/- | 0.40 | 0.25 | 0.50 | 0.25 | 0.13 | 0.38 | 0.25 |
| 15A Average | 1.44<sup>**</sup> | 3.06<sup>**</sup> | np | 2.75<sup>**</sup> | 3.25<sup>**</sup> | np | 6.63 |
| +/- | 0.45 | 0.21 | np | 0.68 | 0.50 | np | 0.52 |
| 6A Average | 1.22<sup>**</sup> | 4.08<sup>**</sup> | 4.50<sup>**</sup> | 4.25<sup>**</sup> | 4.58<sup>**</sup> | 4.15<sup>**</sup> | 8.06 |
| +/- | 0.23 | 0.61 | 0.75 | 0.82 | 0.42 | 0.85 | 0.84 |
| TD10 Average | np | 8.17<sup>ns</sup> | np | 7.50<sup>ns</sup> | np | np | 9.33 |
| +/- | np | 1.18 | np | 0.41 | np | np | 0.47 |
| USDA-ARS Phenotyping | <u>CI5791</u> | <u>HVT02681</u> | <u>HVT02682</u> | <u>HVT02690</u> | <u>HVT02691</u> | <u>HVT02693</u> | <u>Golden Promise</u> |
| 0-1 Average | np | 6.5<sup>*</sup> | np | 5.5<sup>*</sup> | np | np | 7.83 |
| +/- | np | 0.00 | np | 0.00 | np | np | 0.24 |
| 6A Average | np | 3.17<sup>*</sup> | np | 3.5<sup>*</sup> | np | np | 5.67 |
| +/- | np | 0.47 | np | 0.00 | np | np | 0.47 |
| TD10 Average | np | 8.00<sup>ns</sup> | np | 8.33<sup>ns</sup> | np | np | 8.67 |
| +/- | np | 0.24 | np | 0.41 | np | np | 0.62 |

Expression analysis was performed to confirm that the CI5791 *Rpt5* allele was being expressed in transgenic lines. Resistant transgenic lines HVT02681 and HVT02691 were selected to represent individuals generated from callus #16 and callus #19, respectively. Results of the qPCR assay confirmed expression in both lines, with an average normalized expression level of 4.8 for HVT02681 and 7.3 for HVT02691 before pathogen inoculation (0 hours). HVT02681 expression levels were also analyzed 48 hours post inoculation with *Ptt* isolate 6A and showed an average expression level of 5.7. Highly resistant line CI5791 showed similar expression levels prior to pathogen inoculation (4.4), however was induced 48 hours post inoculation to a relative expression level of 21.9 (Figure 5). The lack of induction in the transgenic lines was expected given that the *Mla6* promoter drives moderate constitutive expression (Bettgenhaeuser *et al*., 2021; Halterman *et al*., 2001). The difference in induction of *Rpt5* in CI5791 -vs- the *Rpt5* transgenic lines post pathogen inoculation may explain some of the phenotypic difference observed between the transgenic lines (average 6A disease score = 3.85) and CI5791 (average 6A disease score = 1). Conversely, wildtype Golden Promise displayed no detectable expression at 0 hours and 48 hours post inoculation, which was anticipated as the qPCR primers were designed to specifically target the *Rpt5.CI5791* allele (Figure 5). These findings support the role of *Rpt5* as a broad spectrum *Ptt* resistance gene.

**Figure 5.**
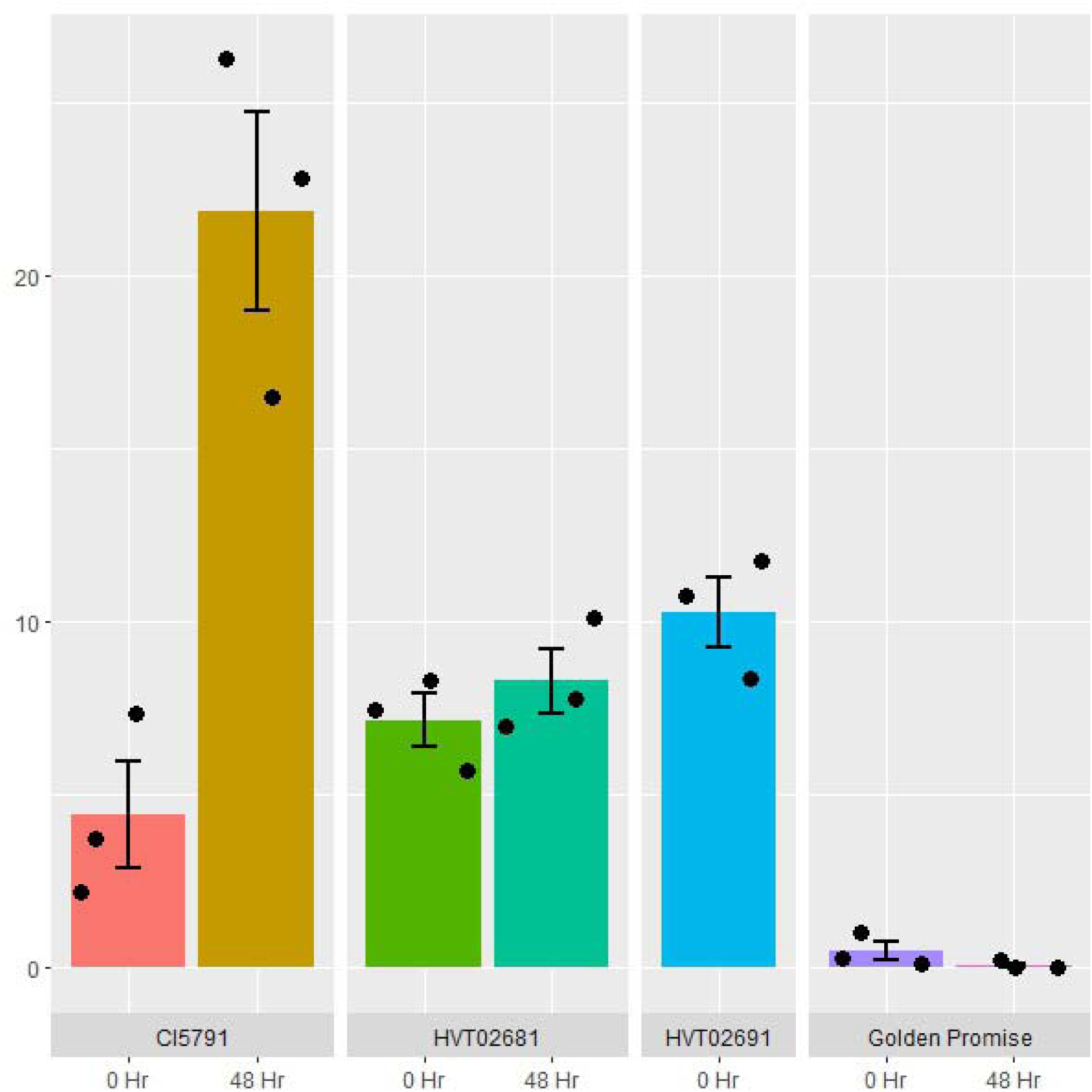
*Rpt5.CI5791* is induced in WT CI5791 and constitutively expressed in transgenic lines. Relative expression levels of *Rpt5.CI5791* in resistant parent, CI5791 (left), susceptible transgenic parent, Golden Promise (right), and transgenic lines HVT02681 and HVT02691. *Rpt5* is strongly induced (∼5-fold) 48 hours following *Ptt* inoculation in CI5791 which may contribute to the robust early resistance observed in this line. Error bars represent +/- standard error of the mean across replications. The expression was normalized to reference gene *HvSnoR14*.

Notable differences in seed morphology, germination rate, growth rate, plant leaf color, and overall plant architecture were observed between transgenic lines and the wildtype background line Golden Promise (shown in Figure S2). To ensure these plants were truly Golden Promise transgenics and not the result of incidental seed admixture, we genotyped transgenic lines HVT02681, HVT02690, HVT02691 and HVT02693 and parental line Golden Promise with the iSelect barley 50K genotyping chip. Genotyping was performed at the USDA North Central Small Grains Genotyping Lab in Fargo, ND (Bayer *et al*., 2017). Analysis of SNPs revealed ∼99.98% identity between all transgenic lines and Golden Promise, confirming that these plants were genuine transgenics of Golden Promise origin (Dataset S2). We hypothesize that the phenotypic differences observed between these transgenic lines and Golden Promise are the result of constitutive expression of *Rpt5.CI5791* driven by the *Mla6* promoter (Halterman *et al*., 2001). Previous RNA seq studies have revealed tightly controlled expression of *Rpt5*, with early upregulation during germination and early root and shoot development, and very little to no expression at later developmental stages (Coulter *et al*., 2022; Liew *et al*., 2020; Mascher *et al*., 2017). This data, combined with our findings on the role of *Rpt5* in *Ptt* resistance, has led us to hypothesize that *Rpt5* has evolved pleiotropic effects in plant growth and development in addition to disease resistance. However, further studies are necessary to elucidate if *Rpt5* has a role in plant growth and development or if the morphological differences are just due to aberrant expression of *Rpt5*.

## Discussion

Fungal diseases present major threats to agricultural production, and climate change will exacerbate these issues as temperatures rise and rainfall patterns are altered (Shiferaw *et al*., 2011; Tubiello *et al*., 2007). Modelling studies have predicted that the supply of malt-quality barley will be impacted by climate change, threatening brewing and distilling industries globally (Xie *et al*., 2018). Moreover, studies predicting future rainfall patterns suggest that many barley growing regions will experience an increase in days conducive to net blotch infection (Launay *et al*., 2014). These additional days of potential disease development are likely to occur early in the growing season, increasing cycles of secondary infections that could lead to more destructive and frequent net blotch epidemics. While fungicide application can limit the impact of net blotch, emerging fungicide resistance (summarized in Effertz *et al*., (2021) and the rising cost of fuel and fungicide makes the deployment of genetic resistance essential for NFNB control in barley (Effertz *et al*., 2021). While multiple net blotch resistance/susceptibility loci have been genetically characterized, no resistance or susceptibility genes had been previously identified and validated.

Research into the genetic interactions between *Ptt* and barley have been ongoing for nearly 100 years (Geschele, 1928). Over 30 independent loci across all seven barley chromosomes have been genetically characterized as resistance or susceptibility loci contributing to *P. teres* disease outcome. The majority of these loci are specific to *Ptt* or have roles in both *Ptt-* and *Ptm-*mediated resistances (summarized in Clare *et al*. (2020). The *Rpt5/Spt1* locus on chromosome 6H has been reported as a robust source of resistance in a variety of biparental and genome wide association studies (Adhikari *et al*., 2020; Amezrou *et al*., 2018; Daba *et al*., 2019; Emebiri *et al*., 2005; Graner *et al*., 1996; Grewal *et al*., 2008; Gyawali *et al*., 2019; Khan and Boyd, 1969; Ma *et al*., 2004; Manninen *et al*., 2000, 2006; O’Boyle *et al*., 2014; Raman *et al*., 2003; Vatter *et al*., 2017; Wang *et al*., 2015; Wonneberger *et al*., 2017). The locus also contains multiple isolate-specific susceptibility genes/alleles identified in several diverse biparental populations (Abu Qamar *et al*., 2008; Richards *et al*., 2017, 2016; Steffenson *et al*., 1996). *Rpt5* alleles from the Ethiopian landraces CI5791 and CI9819 confer the most effective and broadest resistance known in the barley-*Ptt* pathosystem (Koladia *et al*., 2017; Manninen *et al*., 2000, 2006; Robinson and Jalli, 1996).

In this study, we identified and validated *Rpt5*, a gene encoding a receptor-like protein (RLP) with substantial diversity across barley genotypes (Figure 2). Two independent transgenic lines in the Golden Promise genetic background carrying the *Rpt5* allele from the highly resistant landrace CI5791 (*Rpt5.CI5791)* showed a strong shift towards resistance when challenged with the *Rpt5* avirulent *Ptt* isolates 6A, 15A, and 0-1. Expression of the CI5791 *Rpt5* allele in the Golden Promise transgenic lines driven by the *Mla6* promoter resulted in moderate levels of constitutive expression, which conferred less effective resistance than the CI5791 allele with its native promotor that was induced upon pathogen recognition (Bettgenhaeuser *et al*., 2021). Wildtype CI5791 showed similar *Rpt5* expression levels to the transgenic lines prior to inoculation, however *Rpt5* was induced ∼5-fold in CI5791 48 hours after pathogen challenge. We hypothesize the difference in *Rpt5* induction may explain the differential reaction types between CI5791 and the Golden Promise transgenic lines suggesting that the amplitude of *Rpt5* expression may impact *Rpt5*-mediated resistance. Phenotyping of transgenic lines harboring multiple copies of *Rpt5.CI5791* with isolate 6A, 15A, and 0-1 revealed a reaction more similar to wildtype Golden Promise, and expression analysis revealed these lines showed lower transgene expression than single copy lines (Figure S3). We predict these lower expression levels are due to the multiple copies inducing homology dependent gene silencing (Meyer and Saedler, 1996; Rajeevkumar *et al*., 2015), leading to decreased transgene expression and a susceptible outcome when challenged with *Ptt*. However, the transgenic lines containing the Mla6:*Rcg1.CI5791* single copy insertions displayed a significant shift from susceptibility to resistance as compared with WT Golden Promise, showing that the RLP is *Rpt5*.

Plants and fungal pathogens co-evolve in host-pathogen molecular arms races with the pathogen evolving effectors to overcome resistance and the host evolving the ability to recognize and mount an effective defense response (Jones and Dangl, 2006). As research suggests that alterations to environmental conditions may increase the evolutionary rate of fungal pathogens (Möller and Stukenbrock, 2017; Velásquez *et al*., 2018) it becomes even more important to understand the evolutionary mechanisms acting at agronomically relevant resistance/susceptibility loci to ensure that effective resistance is intelligently deployed with durability in mind. Pathogens’ rapid evolution towards virulence applies selective pressure on the plant immunity mechanisms to evolve effective means to suppress colonization (Anderson *et al*., 2010; Cesari, 2018; Fei *et al*., 2016; Jones and Dangl, 2006). Plants, in particular the cereal grains, harbor large, highly plastic, and repeat-rich genomes (Jayakodi *et al*., 2020; Kapustová *et al*., 2019; Williams *et al*., 1999). These repeat rich regions are largely gene-poor and comprised of transposable elements and other DNA sequences of unknown function. Interestingly, these regions are more likely to experience large scale genome rearrangements, such as inversions or gene conversion (Fang *et al*., 2014; Huang and Rieseberg, 2020; Jayakodi *et al*., 2020; Wang and Paterson, 2011). The *Rpt5*/*Spt1* locus is near the centromere of barley chromosome 6H, encompassing a gene poor, transposable element rich region that appears to contain a conserved mechanism that directs non-canonical, double recombination events flanking and delimiting the *Rpt5* locus. Approximately 1% of CI5791 x Tifang F_2_ high-resolution mapping individuals and 3-4% of CI5791 x Tifang RIL and Steptoe x Morex DH individuals screened in our and previous mapping efforts contained double recombination events flanking and delimiting the *Rpt5/Spt1* locus. These double recombination events appear to occur at a conserved location that maintains the *Rpt5/Spt1* locus as a continuous block during meiosis and transmission. We hypothesize that multiple genes within the *Rpt5/Spt1* locus may function in barley-*Ptt* host-pathogen genetic interactions that mediate resistance/susceptibility, and that these double recombination events may represent a conserved mechanism to transfer this important locus generationally as a single block. The presence of highly diverged CI5791 copies of the RLK-encoding genes *Rcg2* and *Rcg3* present at this locus suggest they may also play a role in either *Rpt5*-mediated resistance or *Spt1*-mediated susceptibility, although all attempts to date to amplify the CI5791 alleles of these genes have failed. Further work is required to explore whether/how these genes function in this pathosystem. The lack of easily deployable genetic markers saturating gene poor regions of barley chromosomes increases the challenge of mapping and characterizing non-canonical recombination events as we observed in this study. Further genetic and functional studies are necessary to elucidate the mechanism underlying these double recombination events, which will likely require the development of novel methods to uncover and dissect the processes underlying these genetic phenomena.

*Rpt5*-mediated resistance derived from CI5791 is effective against nearly all global *Ptt* isolates. However, this resistance has been overcome by pathogen populations which had a high level of selection pressure applied through barley monoculture comprised of lines containing the *Rpt5* resistance. One such population from Morocco evolved virulence on *Rpt5* following the deployment of barley cultivar Rabat071, which is known to harbor the CI5791 allele of *Rpt5* (Richards et al., 2024; Li et al., 2024; Taibi *et al*., 2016) and was grown on a large scale over a long period. It is hypothesized that diversifying pressure from several decades of Rabat071 cultivation resulted in the pathogen evolving an effector present on *Ptt* chromosome 1 that enabled the isolates to overcome *Rpt5*-mediated resistance (Li et al., 2024). Now that *Rpt5* has been identified, additional functional studies are underway to determine the mechanism of *Rpt5*-resistance as well as the evolution of *Rpt5*-virulence.

Our group recently published the identification of *HvWRKY6*, a transcription factor conserved across barley genotypes that is required for *Rpt5*-mediated resistance (Tamang *et al*., 2021). This gene was discovered through the screening of ∼10,000 gamma irradiated mutant plants in the CI5791 background. Interestingly, no *Rpt5* mutant was identified in this screen or further screening efforts of ∼20,000 additional CI5791 mutants, leading us to hypothesize that perhaps *Rpt5* knockouts are lethal. Previous reports showed that *Rpt5* was expressed during seed germination and early root and shoot development, however very little expression is observed in the developing inflorescence and in the grain post-anthesis (Coulter *et al*., 2022; Liew *et al*., 2020; Mascher *et al*., 2017). We observed that transgenic lines HVT02681, HVT02690, and HVT02691 display an altered seed and spike phenotype (Figure S2), likely due to the constitutive expression of *Rpt5.CI5791* driven by the *Mla6* promoter during inflorescence and grain development. These lines also display reduced germination rates compared to Golden Promise. Taken together these findings suggest a potential role for *Rpt5* in early plant development in addition to disease resistance.

The research outlined in this study and our previous research on *HvWRKY6* provide two key pieces of information towards understanding the mechanisms of broad *Rpt5*-mediated resistance and *Spt1*-mediated genotype specific susceptibility. While the functional characterization of resistance in pathosystems such as rice-rice blast and wheat-wheat stem rust are being investigated, functional characterization studies in the barley-net blotch pathosystem were not possible until net blotch resistance or susceptibility genes had been identified. Identifying the genes underlying the *Rpt5/Spt1* locus provides the tools to begin this characterization, as the locus is involved in multiple interactions that mediate resistant and susceptible outcomes via gene-for-gene dominant resistance as well as multiple genotype specific inverse gene-for-gene susceptibility interactions. Functional characterization studies are currently underway to determine the host and pathogen-derived factors that influence *Rpt5*-mediated resistance and *Spt1*-mediated susceptibility.

## Materials and Methods

### *Pyrenophora teres* f. *teres* inoculations and phenotyping

*Ptt* isolates were obtained from Dr. Brian Steffenson and Dr. Tim Friesen, with isolates 6A and 15A originating from California, USA, isolate TD10 originating in Montana, USA, isolate LDN89-19 originating in North Dakota, and isolate 0-1 originating from Ontario, Canada. *Ptt* culturing, inoculations, and phenotyping for high-resolution mapping and subsequent analysis of transgenic lines was carried out as described in Friesen *et al*. (2006). Briefly, barley plants were grown to the two-leaf stage (∼14 days) before being spray inoculated with a *Ptt* spore suspension at ∼2,000 spores/ml until runoff. Plants were then kept at 100% humidity in a mist chamber for 22-24 hours before being returned to growth chamber conditions. Plants were rated 7 days post inoculation using the Tekauz 1-10 *Ptt* rating scale (Tekauz, 1985).. All transgenic lines resulting from callus #16 (HVT02681, HVT02682), callus #19 (HVT02690, HVT02691, HVT02693), which had shown significant shifts towards resistance, along with multi-copy transgenics HVT02680 and HVT02686 were phenotyped with isolates 6A and 0-1. Due to low seed number and poor germination, only HVT02681, HVT02690, HVT02691, HVT02680, and HVT02686 were phenotyped with isolate 15A, and HVT02681, HVT02690, HVT02680, and HVT02686 were phenotyped with isolate TD10. Phenotyping of each transgenic line was performed in triplicate (three plants per technical rep) with at least three technical reps per isolate.

### High-resolution mapping

Primers targeting genic regions of the Morex V1 assembly were manually designed to saturate the region between previously published flanking markers 11_20946 and SCRI_RS_187343. Approximately 1,000 bp amplicons were amplified from parental lines CI5791 and Tifang and Sanger sequenced (Genscript) to determine appropriate targets for single nucleotide polymorphism (SNP) markers. Following SNP identification new markers were designed for sequencing on the Ion Torrent platform, with an amplicon size of ∼200bp and Ion Torrent adapter sequences added to the 5’ ends of each forward and reverse primer. Saturating and flanking markers were pooled and used to screen 960 CI5791 x Tifang F_2_ individuals representing 1920 recombinant gametes. F_2_ individuals were planted in 98-rack conetainers and primary leaves were sampled for DNA extraction. All F_2_ individuals were phenotyped with *Ptt* isolate 0-1 as described above. Genomic DNA extraction, library preparation, Ion Torrent sequencing, and bioinformatic analysis were performed as described in Sharma Poudel *et al*. (2018). After the completion of genotyping and phenotyping individuals harboring recombination between flanking markers 11_20946 and SCRI_RS_187343 were transplanted and raised to maturity for the generation of immortal critical recombinants. Additional STS markers were developed targeting three candidate genes within the delimited *Rpt5* region, allowing for rapid confirmation of genotype in this region without necessitating a full sequencing run. These markers were also used to screen individuals from the CI5791 x Tifang RIL population and the Steptoe x Morex DH population (Chen and Hayes, 1989; V. M. Koladia *et al*., 2017). Results from these additional population screens are reported in Dataset S1.

### CI5791 PacBio sequencing, Allele analysis, and candidate gene selection

Initial analysis of the gene space delimited by our high-resolution mapping efforts revealed several potential candidate genes. Primers designed to amplify potential candidate genes failed to amplify most CI5791 alleles, leading us to employ low coverage PacBio sequencing of CI5791 for allele mining. Seven-day old seedlings were grown in normal greenhouse conditions prior to a 24-hour dark treatment that immediately preceded tissue collection. Sixty grams of leaf tissue was collected and immediately flash frozen in liquid nitrogen before being shipped on dry ice to the Arizona Genomics Institute (AGI, University of Arizona) for high-molecular weight DNA extraction and PacBio Sequel II (PacBio, https://www.pacb.com) sequencing. A single continuous long read (CLR) run was performed at AGI, yielding 139 GB of data and approximately 28X coverage of the CI5791 genome. Reads were assembled using Canu (v2.1.1) with default settings (genome size = 4.1 Gb) (Koren *et al*., 2017). Although the integrity of the *Rpt5* region was incomplete in this assembly, it proved useful for CI5791 allele mining. Susceptible parent Tifang was found to possess a 100% identical *Rcg1* allele to Morex, as well as identical SNPs within the GBS markers in and surrounding the locus. These findings guided our decision to use the Morex V3 assembly for allele comparisons within the *Rpt5* delimited region. Allele comparisons were performed within Geneious v2022.2.2 using Morex gene sequences in BLAST searches against the CI5791 assembly. Genes containing SNPs between Morex and CI5791 were analyzed for potential amino acid substitutions. This analysis left ten genes with non-conservative amino acid differences, including a duplicated receptor-like protein (*HORVU.MOREX.r3.6HG0595020*(*Rcg1*) and *HORVU.MOREX.r3.6HG0594600*, 71% nucleotide identity) and a duplicated receptor-like kinase (*HORVU.MOREX.r3.6HG0595000 (Rcg2)* and *HORVU.MOREX.r3.6HG0594730 (Rcg3)*, 62% nucleotide identity).

### Cloning and stable transformation of *Rcg1.CI5791* in “Golden Promise”

Identification of *Rcg1* as a potential candidate gene occurred prior to our PacBio sequencing effort. Due to the extreme diversity present at the 5’ end of *HORVU.MOREX.r3.6HG0595020 (Rcg1)*, we chose to utilize a genome walking strategy to obtain the full-length allele from CI5791, as PCR primers designed from the Morex assembly failed to amplify the 5’ region of the CI5791 allele of this gene. Genome walking was performed using a modified Genome Walker 2.0 kit (Clonetech (now Takara Bio) https://www.takarabio.com/). CI5791 genomic DNA was overnight digested with four blunt end restriction enzymes, *Dra*I, *Eco*RV, *Nae*I, and *Pvu*II, purified, and ligated to GenomeWalker adaptors to generate four genome walking libraries. Nested PCR was then performed using several gene specific primers (Table S2) designed in the more conserved LRR region of *Rcg1* combined with a primer targeting the GenomeWalker AP1 adaptor sequence. Primary PCR products were diluted 1:50 and used as the templates for secondary PCR, which utilized nested gene specific primers paired with a primer targeting the AP2 adaptor sequence. Secondary PCR products were run out on 1% agarose gel, positive bands were excised, and the DNA was purified using the Wizard SV Gel Clean-Up System (Promega, https://www.promega.com). Purified products were sent for bi-directional Sanger sequencing through Genscript.

After obtaining the full-length *Rcg1.CI5791* allele, cloning primers were designed with *Hin*dIII (F) and *Spe*I (R) enzyme cut sites in the forward and reverse primers. The full-length *Rpt5* allele was amplified from CI5791 cDNA. The amplicon and target vector, pBract202, were digested with the restriction enzymes *Hind*III and *Spe*I (New England Biolabs, https://www.neb.com). Ligations were performed using T4 ligase (NEB). The ligation reactions were transformed into Top10 *E. coli* cells (Invitrogen, https://www.thermofisher.com/us/en/home/brands/invitrogen.html) and plated on LB plates containing 50 µg/ml neomycin for selection. Colonies were screened initially by PCR and positive colonies were selected for plasmid extraction and Sanger sequencing to confirm in frame cloning of the allele. *Mla6* promoter and terminator sequences were cloned into the vector up and downstream of the *Rcg1.CI5791* allele, as we were unable to obtain the native promoter and terminator from CI5791 via genome walking. The *Rcg1.CI5791*-pBract202 construct was transformed at The Sainsbury Lab (Norwich, England, UK) using the barley accession Golden Promise. An annotated vector map of the construct used for transformation has been deposited in GenBank under accession OP868846.

*Agrobacterium*-mediated transformation was performed as described in Bartlett *et al*. (2008). Briefly, Golden Promise plants were grown in growth chamber conditions until immature embryos were ∼1.5 mm in diameter. Immature embryos were then isolated and incubated with a fresh culture of *Agrobacterium* Agl1 harboring the *Rcg1.CI5791*-pBract202 construct on fresh callus induction (CI) media without antibiotics. After three days of incubation on CI media, the embryos were transferred to a selectable CI media containing Timentin and Hygromycin and incubated in the dark for six weeks at 24°C, with fresh media being provided every two weeks. After six weeks of callus induction, the calli were transferred to transition medium and cultured in the light at 24°C for two weeks before transferring to regeneration media until roots and shoots developed. The plantlets were then transferred to rooting tubes until the shoots had reached to top of the rooting tubes at which point, they were transferred to soil and grown to maturity. Transgene copy number of transgenic plants was confirmed via qPCR using the *hyg* selectable marker (iDNA Genetics, Norwich, UK) before the seeds were sent to WSU for phenotyping of the transgenic lines.

### Pooled primer amplification and sequencing of *Rcg1* alleles from diverse barley lines

Given the high level of sequence diversity present in the 5’ region of *Rcg1* across alleles, we employed a pooled primer PCR system to amplify and sequence alleles from a selection of cultivars and net blotch resistant lines from the world barley core collection (Richards *et al*., 2017). Primers utilized are outlined in Table S2. Primer pools were used to amplify *Rcg1* alleles using a thermal cycler program of an initial melt at 95°C for 3 minutes, 35 cycles of 95°C for 30 seconds, 50°C anneal for 30 seconds, 72°C extension for two minutes, followed by a five-minute final extension at 72°C and a final hold at 4°C. Products were run out on a 1% agarose gel and bands of 1200-1400 bp were excised and subsequently gel extracted with the Wizard SV Gel Cleanup Kit (Promega, https://www.promega.com). Following gel extraction, products were fragmented using NEBNext® dsDNA Fragmentase® (New England Biolabs, https://www.neb.com) to approximately ∼200 bp fragments, barcoded, and sequenced on the Ion Torrent PGM as described above. Products were *de novo* assembled and aligned to known *Rcg1* alleles from barley lines Rika, Kombar, Morex, and CI5791 using CLC Genomics workbench. Assembled contigs have been deposited into GenBank under BioProject PRJNA907319, accession KFXA00000000.

### Analysis of *Rpt5* genetic diversity and protein structure

The coding sequence of the *Rcg1* gene family was aligned using Geneious translation alignment that uses MUSCLE (v3.8.425) using default parameters. The maximum likelihood phylogenetic tree was generated using RAxML (v8.2.12) with the general time reversible CAT model (GTRCAT). Phylogenetic tree visualization was performed with iTOL (https://itol.embl.de). Protein structure prediction was performed using the AlphaFold2 Colab implementation (simplified version of AlphaFold2 v2.2.4; https://colab.research.google.com/github/deepmind/alphafold/blob/main/notebooks/AlphaFold.ipynb). Protein structure visualization was performed with PyMOL (v2.5.2).

### Transgene expression analysis

Transgenic lines HVT02681, and HVT02691 along with parental line Golden Promise were grown in growth chamber conditions and inoculated with *Ptt* isolate 6A as described above. One-to-two-inch samples of secondary leaf tissue were collected, and flash frozen in liquid nitrogen from all lines in triplicate prior to inoculation (0 hr), and 48 hours post inoculation. RNA was extracted from tissue using the Qiagen RNeasy mini kit (Qiagen) and cDNA was synthesized using the GoScript reverse transcription system (Promega, https://www.promega.com). Primers designed to specifically target the *Rpt5.CI5791* allele were developed to determine the level of transgene expression and native expression in CI5791. qPCR was conducted using SsoAdvanced™ SYBR® Green Supermix (Bio-Rad) on the CFX384 Real-time thermal cycler (Bio-Rad) with the cycling parameters of 95°C for 30 s followed by 40 cycles of 95°C for 15 s and 58°C for 30 s; 65°C for 30 s; and 60 cycles of temperature increasing from 60 to 95°C with fluorescence readings acquired at 0.5°C increments per cycle. Three technical replications were performed for each biological replication, at least three biological replications were utilized for each treatment. The barley gene *HvSnoR14* was used for reference to normalize *Rpt5.CI5791* expression (Ferdous *et al*., 2015; Tamang *et al*., 2021). Relative expression levels were determined using the ΔΔCT method on the Bio-Rad CFX Manager 3.1 software.

## Acknowledgments

The authors would like thank Tom Gross and Pat Gross for assistance with Ion Torrent sequencing and greenhouse support during the high-resolution mapping phase of this project, Dave Kudrna for advice and assistance with PacBio Sequencing, Inmaculada Hernández-Pinzón with molecular cloning of *Rpt5*, Matthew Smoker and Jodi Taylor for barley transformation, and Phon Green with greenhouse assistance of transgenic plants. The research presented in this manuscript was supported by the USDA National Institute of Food and Agriculture Hatch project 1014919, Crop Improvement and Sustainable Production Systems (WSU reference 00011), in addition to funding from the United States Department of Agriculture (Award #2018-67014-28491, CRIS #5062-21220-025-000D), the United States National Science Foundation (Award #1759030), and the Gatsby Charitable Foundation.

