## Supplementary Figures and Tables for "*Rpt5* encodes a receptor-like protein that provides broad and effective net form net blotch (*Pyrenophora teres* f. *teres*) resistance in barley"

**This PDF file includes:**

Figures S1 to S3
Tables S1 to S3

**Other supporting materials for this manuscript include the following:**

Datasets S1 to S2

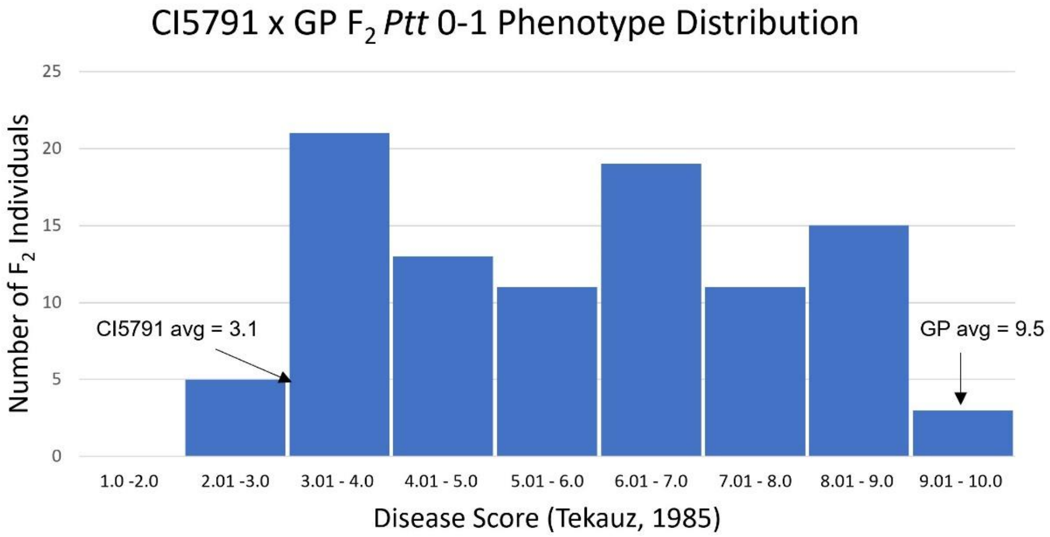


**Figure S1.** Disease score distribution of CI5791 x Golden Promise (GP) F2 individuals (n=98) phenotyped with *Ptt* isolate 0-1. Average parental scores for this phenotyping trial were 3.1 for CI5791 and 9.5 for Golden Promise. CI5791 exhibited a higher disease rating than typically observed during this assay, potentially due to a spore concentration higher than 2,000/mL. Distribution of scores suggests that factors other than a single dominant resistance gene (*Rpt5*) influence disease outcome in this cross.

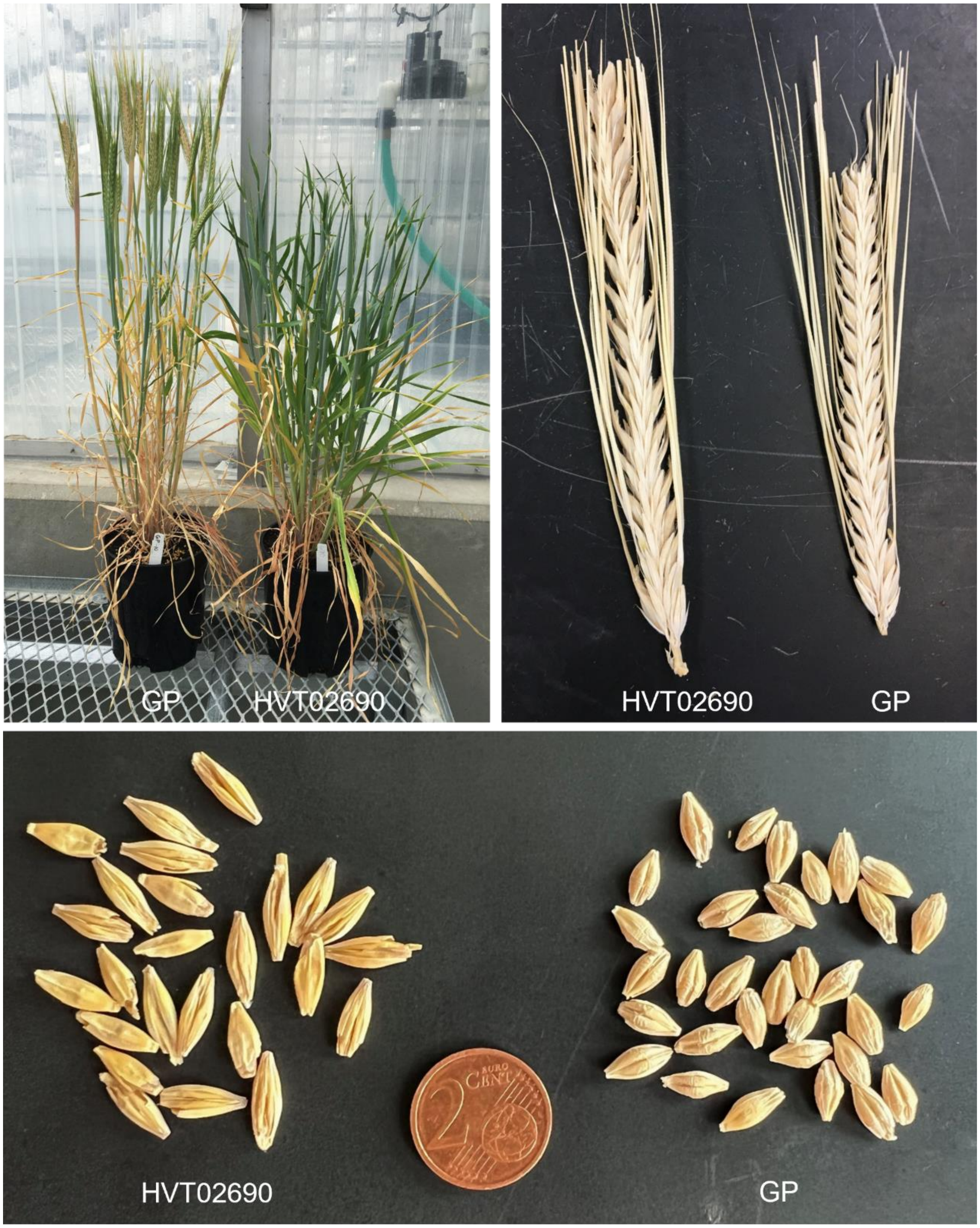


**Figure S2.** Golden Promise single-copy transgenic line (HVT02690) expressing *Rpt5.CI5791* displayed altered plant architecture and development (left), head morphology (right), and seed shape (below) as compared to wildtype (GP). We hypothesize these differences are due to constitutive expression of *Rpt5.CI5791* driven by the *Mla6* promoter in our transgenic construct.

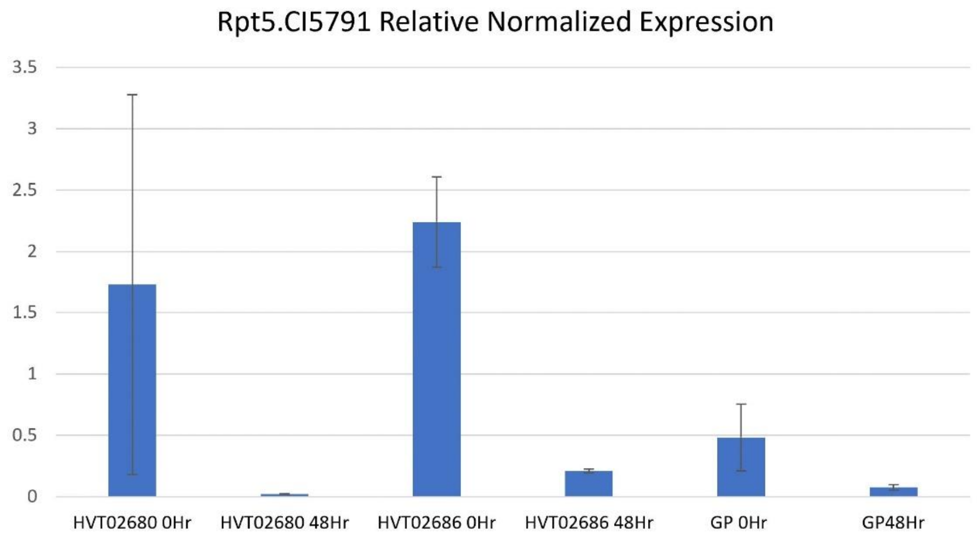


| | HVT02680 (4 copies) | HVT02686 (3 copies) | Golden Promise |
|---|---|---|---|
| *Ptt*6A | 6.333 +/- 0.471 | 5.417 +/- 0.425 | 6.500 +/- 0.707 |
| *Ptt*15A | 6.667 +/- 1.247 | 6.5 +/- 0.500 | 7.25 +/- 0.433 |
| *Ptt*0-1 | 9.5 +/- 0.408 | 9.167 +/- 0.623 | 9.833 +/- 0.236 |

**Figure S3 –** Multi-copy transgenic lines show lower transgene expression and have Ptt phenotypes similar to wildtype Golden Promise. +/- indicates standard deviation across three phenotypic replicates for each isolate.

**Table S1.** Disease reactions and *Rcg1* haplotype of barley genotypes screened in this study.

| Barley Genotype | *Pyrenophora teres* f. *teres* disease rating | | | | | *Rcg1* Haplotype |
|---|---|---|---|---|---|---|
| | 6A | 15A | LDN89-19 | TD10 | 0-1 | |
| CI5791 | 1.000 | 1.000 | 1.000 | 1.250 | 1.300 | CI5791 |
| Morex* | 2.700 | 2.200 | 7.000 | 7.750 | 7.750 | Morex |
| Tifang | 1.750 | 1.500 | 7.100 | 7.750 | 8.100 | Morex |
| Rika | 8.700 | 1.700 | NA | 3.500 | 5.200 | Rika |
| Kombar | 2.250 | 7.500 | NA | 6.000 | 9.330 | Kombar |
| WBD209 | NA | NA | 1.500 | 3.500 | 3.000 | CI5791 |
| ND Genesis | 2.000 | 1.333 | 1.333 | 2.333 | 2.333 | CI5791 |
| Heartland | 1.333 | 2.333 | 2.000 | 2.000 | 2.000 | CI5791 |
| Conlon | 2.000 | 2.000 | 2.000 | 2.333 | 2.000 | CI5791 |
| Golden Promise* | 7.000 | 7.500 | NA | 9.333 | 9.250 | Morex |
| Rabat071 | 1.333 | 1.333 | 2.000 | 1.667 | 1.667 | CI5791 |
| CI9819 | 2.000 | 1.333 | NA | 2.000 | 2.050 | CI5791 |
| CI11458 | 7.000 | 2.000 | NA | 5.000 | NA | Rika |
| Hazera | 5.333 | 1.500 | NA | 1.500 | NA | Kombar |
| PI189770 | 8.000 | 1.800 | NA | 7.250 | NA | Morex |
| Hector | 7.500 | 7.750 | NA | 8.750 | NA | Morex |
| NDB112 | 1.250 | 1.000 | NA | 7.000 | NA | Morex |
| HOR13942* | 2.000 | 5.000 | NA | NA | 4.333 | Other |
| HOR3081* | 3.500 | 3.333 | NA | NA | 6.333 | Rika |
| HOR3365* | 2.000 | 2.667 | NA | NA | 5.500 | Other |
| HOR13821* | 2.500 | 1.167 | NA | NA | 6.333 | Kombar |
| Bowman | 5.500 | 2.167 | NA | NA | 7.667 | Morex |
| MXB468 | 6.667 | 6.000 | NA | NA | 10.000 | Unknown |
| HOR17287 (Igri)* | 7.667 | 4.333 | NA | NA | 10.000 | Other |
| HOR21599* | 1.833 | 4.667 | NA | NA | 7.000 | Morex |
| HOR9043* | 3.500 | 2.000 | NA | NA | 5.333 | Kombar |
| HOR8148* | NA | 1.000 | NA | NA | 8.667 | Kombar |
| HOR7552* | 8.500 | 4.333 | NA | NA | 8.500 | Other |
| HOR10350* | 3.167 | 3.333 | NA | NA | 4.667 | Other |
| HOR 18579 (Akashinriki)* | NA | 5.000 | NA | NA | 6.000 | Other |
| OUN333* | 6.000 | 2.333 | NA | NA | 9.333 | Other |
| Barke* | NA | 2.000 | NA | NA | 5.000 | Other |
| Hockett* | 2.417 | 2.083 | NA | NA | 5.833 | Morex |
| PI564598 | 1.667 | 1.500 | 1.833 | NA | NA | CI5791 |
| PI564637 | 1.000 | 1.500 | 1.000 | NA | NA | CI5791 |
| PI573611 | 2.167 | 2.167 | 1.833 | NA | NA | CI5791 |
| PI584971 | 1.000 | 1.833 | 1.167 | NA | NA | CI5791 |
| PI226639 | 1.833 | 2.333 | 2.333 | NA | NA | CI5791 |
| PI237581 | 2.167 | 2.500 | 1.500 | NA | NA | Unknown |
| PI264208 | 1.000 | 2.167 | 1.167 | NA | NA | Unknown |
| PI264209 | 1.333 | 1.500 | 1.833 | NA | NA | CI5791 |
| PI295958 | 1.667 | 1.667 | 2.000 | NA | NA | Unknown |
| PI296192 | 1.000 | 1.667 | 1.333 | NA | NA | CI5791 |
| PI319867 | 1.167 | 1.333 | 1.500 | NA | NA | Unknown |
| PI330398 | 1.500 | 2.333 | 1.833 | NA | NA | Unknown |
| PI344936 | 1.167 | 2.333 | 1.833 | NA | NA | Rika |

| | | | | | | |
|---|---|---|---|---|---|---|
| PI356580 | 1.250 | 1.500 | 1.500 | NA | NA | Unknown |
| CIho5057 | 1.333 | 2.333 | 1.667 | NA | NA | CI5791 |
| CIho7492 | 1.500 | 1.333 | 1.167 | NA | NA | Unknown |
| CIho7504 | 1.333 | 1.833 | 1.833 | NA | NA | CI5791 |
| CIho11533 | 1.000 | 1.833 | 1.667 | NA | NA | Unknown |
| CIho14260 | 1.167 | 1.500 | 1.000 | NA | NA | Unknown |
| CIho14286 | 1.167 | 2.000 | 1.833 | NA | NA | CI5791 |
| CIho14880 | 1.333 | 1.333 | 1.667 | NA | NA | CI5791 |
| CIho15364 | 1.333 | 1.500 | 1.333 | NA | NA | CI5791 |
| PI605699 | 1.667 | 1.500 | 1.667 | NA | NA | CI5791 |
| PI606305 | 1.000 | 2.333 | 1.000 | NA | NA | CI5791 |
| PI608667 | 1.500 | 2.167 | 1.333 | NA | NA | Unknown |
| PI611472 | 1.000 | 1.333 | 1.167 | NA | NA | Rika |
| PI611534 | 1.000 | 1.000 | 1.167 | NA | NA | Unknown |
| PI611599 | 1.000 | 1.500 | 1.500 | NA | NA | Rika |
| PI620640 | 1.167 | 1.000 | 1.000 | NA | NA | CI5791 |
| PI636084 | 2.000 | 2.000 | 2.333 | NA | NA | Unknown |
| PI643244 | 1.667 | 1.333 | 1.000 | NA | NA | CI5791 |
| PI643310 | 1.000 | 1.667 | 1.000 | NA | NA | CI5791 |
| PI643314 | 1.000 | 1.000 | 1.000 | NA | NA | CI5791 |
| PI643364 | 2.500 | 1.500 | 1.333 | NA | NA | CI5791 |
| PI26179 | 1.000 | 1.500 | 1.333 | NA | NA | CI5791 |
| PI28624 | 1.667 | 2.167 | 1.333 | NA | NA | CI5791 |
| PI67381 | 1.000 | 1.667 | 1.000 | NA | NA | Rika |
| PI151795 | 1.000 | 1.500 | 1.333 | NA | NA | CI5791 |
| PI159126 | 1.333 | 1.333 | 1.000 | NA | NA | CI5791 |
| PI168328 | 1.667 | 2.167 | 1.167 | NA | NA | CI5791 |
| PI182681 | 1.750 | 1.000 | 2.000 | NA | NA | CI5791 |
| PI190786 | 1.167 | 2.500 | 1.667 | NA | NA | Rika |
| PI221307 | 1.667 | 1.500 | 1.667 | NA | NA | CI5791 |
| PI223446 | 1.333 | 2.167 | 2.000 | NA | NA | CI5791 |
| PI386601 | 1.667 | 2.333 | 1.667 | NA | NA | CI5791 |
| PI386993 | 1.333 | 1.167 | 1.167 | NA | NA | CI5791 |
| PI422232 | 1.000 | 1.833 | 1.500 | NA | NA | CI5791 |
| PI438585 | 1.500 | 2.250 | 2.167 | NA | NA | CI5791 |
| PI447260 | 1.333 | 1.833 | 1.333 | NA | NA | CI5791 |

*Indicates pangenome lines.

**Table S2.** Primers used in this study

| Primer Name | Sequence (5' – 3') | Application |
| --- | --- | --- |
| *Rcg1*_Var1-F* | ACACTGACGACATGGTTCTACAACCAGCGATGACCAGTTTCCGCGAT | *Rcg1* Variable Region Sequencing |
| *Rcg1*_Var1-R* | TACGGTAGCAGAGACTTGGTCTGCATCTGTTTCCCTAGCAATGACTC | *Rcg1* Variable Region Sequencing |
| *Rcg1*_Var2-F* | ACACTGACGACATGGTTCTACAGCTGATGGAGGAAGAAAAAGTTTAC | *Rcg1* Variable Region Sequencing |
| *Rcg1*_Var2-R* | TACGGTAGCAGAGACTTGGTCTAAGGTTTCGCGTCAAGGGAGATTTG | *Rcg1* Variable Region Sequencing |
| *Rcg1*_Var1C-F* | ACACTGACGACATGGTTCTACATCCAGGCCTCCTCCAACAACTT | *Rcg1* Variable Region Sequencing |
| *Rcg1*_Var1C-R* | TACGGTAGCAGAGACTTGGTCTTTGGCGACGGCGAGGTAGCTCA | *Rcg1* Variable Region Sequencing |
| CI5791_*Rcg1*_EndR | TAAGCAATGACACTAATCCACATAA | CI5791 *Rcg1* Allele Confirmation |
| *Rcg1*_Con-1F | ACGGACCTGCCACCTCTATATATA | *Rcg1* Pooled Primer Amplification |
| *Rcg1*_Con-1R | AGAACCTAGAAAAGAAATGGCCAA | *Rcg1* Pooled Primer Amplification |
| *Rcg1*_D_1F | GGTGTCGAATACTATAGGGTACGCG | *Rcg1* Pooled Primer Amplification |
| *Rcg1*_D_2F | CAGTTTCCGCGATCTTTGCATGATC | *Rcg1* Pooled Primer Amplification |
| *Rcg1*_D_3F | AACACTTACGTTGGAAGTGATCTGGCA | *Rcg1* Pooled Primer Amplification |
| *Rcg1*_D_1R | TCCAGCACGCCGTTCTTGATGAGTG | *Rcg1* Pooled Primer Amplification |
| *Rcg1*_D_2R_T2C | TCCAGCACGCCGTTCTTGACGAGTG | *Rcg1* Pooled Primer Amplification |
| *Rcg1*_Walk_1F | AGCACGCCGTTCTTGATGA | *Rcg1*.CI5791 Primer Walking |
| *Rcg1*_Walk_2F | ACAGCGTGAGGTTGGAGAGGT | *Rcg1*.CI5791 Primer Walking |
| *Rcg1*_Walk_1R | GGCTCTTGTACGTGTTGTGATGCTAC | *Rcg1*.CI5791 Primer Walking |
| *Rcg1*_Walk_2R | GGAATTATGTGGATTAGTGTCATTGC | *Rcg1*.CI5791 Primer Walking |
| *Rcg1*_Walk_3R | CATCGCTGGTGTCGAACTTGTATAG | *Rcg1*.CI5791 Primer Walking |
| *Rcg1*_Walk_4R | TAGGTGAGTCTAGCAAGGCGTGGGA | *Rcg1*.CI5791 Primer Walking |
| *Rcg1.C*_HindIII_F | ATAAAGCTTATGGCCGACCACTTCAAATCTCCC | *Rcg1*.CI5791 cloning into pBract202 |
| *Rcg1.C*_SpeI_F | ATAACTAGTTCACTCATGCAGAGTGGCGTACAC | *Rcg1*.CI5791 cloning into pBract202 |
| *Rcg1.C*_qPCR_F1 | GATTTGCTTGTACCCGCTG | *Rpt5*.CI5791 Expression Analysis |
| *Rcg1.Uni*_qPCR_R1 | TAGCTCAACTTGGAGAAGCC | *Rpt5*.CI5791 Expression Analysis |
| *HvSnoR14*_F | GATGTTTATGTATGATAGTCTGTC | Expression normalization |
| *HvSnoR14*_R | GTCGGGATGTATGCGTGTC | Expression normalization |

*Contains 5' Ion Torrent adapter sequence (highlighted in red)

Table S3 – Morex V3 gene annotations within the *Rpt5* delimited region.

| Morex V3 Gene ID | Morex V3 Annotation | Designation (this study) |
|---|---|---|
| ***HORVU.MOREX.r3.6HG0594570*** | Filament-like plant protein | |
| ***HORVU.MOREX.r3.6HG0594580*** | ADP-ribosylation factor GTPase-activating protein 2 | |
| ***HORVU.MOREX.r3.6HG0594590*** | SPla/RYanodine receptor (SPRY) domain protein | |
| ***HORVU.MOREX.r3.6HG0594600*** | Leucine-rich repeat protein kinase family protein | *Rcg4* |
| ***HORVU.MOREX.r3.6HG0594620*** | LINE-1 reverse transcriptase like | |
| ***HORVU.MOREX.r3.6HG0594720*** | NAC (No Apical Meristem) domain transcriptional regulator superfamily protein | |
| ***HORVU.MOREX.r3.6HG0594730*** | Receptor-like protein kinase | *Rcg3* |
| ***HORVU.MOREX.r3.6HG0594880*** | Tesmin/TSO1-like CXC domain containing protein | |
| ***HORVU.MOREX.r3.6HG0594950*** | NAC (No Apical Meristem) domain transcriptional regulator superfamily protein | |
| ***HORVU.MOREX.r3.6HG0595000*** | Receptor-kinase- putative | *Rcg2* |
| ***HORVU.MOREX.r3.6HG0595020*** | Leucine-rich repeat protein kinase family protein | *Rcg1* |
| ***HORVU.MOREX.r3.6HG0595030*** | methyl-coenzyme M reductase II subunit gamma - putative (DUF3741) | |

**Dataset S1 (separate file).** Genotyping data from the CI5791 x Tifang RIL population and Steptoe x Morex DH population showing the double recombination events detected using saturating STS markers. These events were observed in CxT RIL individuals 4, 5, 68, and 112, and SxM DH individuals 063, 071, 143, 164, and 172.

**Dataset S2 (separate file).** Golden Promise and single-copy transgenic line 50K genotyping results. Tifang and CI9819 genotyping calls are included in columns A and B, respectively, as controls. Columns H-K show the comparisons of calls between HVT02681, HVT02690, HVT02691, and HVT02693 and Golden Promise wildtype. Row 44046 shows the percentage of calls that were identical between GP and transgenic lines, all > 99.97% indicating that they were genuine transgenics of Golden Promise background.
